# A tripartite structure, the complex nuclear receptor element (cNRE), is a *c*is-regulatory module of viral origin required for atrial chamber preferential gene expression

**DOI:** 10.1101/2021.11.18.469087

**Authors:** Luana Nunes Santos, Ângela Maria da Souza Costa, Martin Nikolov, Allysson Coelho Sampaio, Frank E. Stockdale, Gang F Wangø, Hozana Andrade Castillo, Mariana Bortoletto Grizante, Stefanie Dudczig, Michelle Vasconcelos, Nadia Rosenthal, Patricia Regina Jusuf, Paulo de Oliveira, Tatiana Guimarães de Freitas Matos, William Nikovits, Michael Schubert, Mirana Ramialison, José Xavier-Neto

## Abstract

Optimal cardiac function requires appropriate contractile proteins in each heart chamber. Atria require slow myosins to act as variable reservoirs, while ventricles demand fast myosin for swift pumping functions. Hence, myosin is under chamber-biased *cis*-regulatory control to achieve this functional distribution. Failure in proper regulation of myosin genes can lead to severe congenital heart dysfunction. The precise regulatory input leading to cardiac chamber-biased expression remains uncharted. To address this, we computationally and molecularly dissected the quail Slow Myosin Heavy Chain III (SMyHC III) promoter that drives specific gene expression to the atria to uncover the regulatory information leading to chamber expression and understand their evolutionary origins. We show that SMyHC III gene states are autonomously orchestrated by a complex nuclear receptor *cis*-regulatory element (cNRE), a 32- bp sequence with hexanucleotide binding repeats. Using *in vivo* transgenic assays in zebrafish and mouse models, we demonstrate that preferential atrial expression is achieved by the combinatorial regulatory input composed of atrial activation motifs and ventricular repression motifs. Through comparative genomics, we provide evidence that the cNRE emerged from an endogenous viral element, most likely through infection of an ancestral host germline. Our study reveals an evolutionary pathway to cardiac chamber-specific expression.

## Introduction

Vertebrate chambered hearts are efficient pumps organized according to an ancient evolutionary paradigm that divides their circulatory functions into inflow and outflow working modules, the atria, and ventricles, respectively (Simões-Costa et al., 2005). Most current views on cardiac chamber development agree with the principles of epigenesis, in which higher-level structures arise through sequential morphogenetic steps. The gradual three-dimensional organization is accompanied by progressive restriction of cellular fates, from large, early embryonic fields that initially display broad tissue potencies (e.g., epiblast or mesoderm) to terminally differentiated cells, such as most heart cell types. The highly varied cardiac cell fates originate from a combination of mosaic and regulative development processes (José Xavier-Neto et al., 2012). Ultimately, these developmental processes combine clues from the relative position of each group of cells inside the embryo with the information imparted by patterning and migration, which modify the initial relationships between cell progenitors and further restrict fates.

Most efforts in the last twenty years have been dedicated to transpose the blueprints of the above-mentioned cardiac ontogenetic events from the four-dimension arena into the overlapping chemical (i.e., signaling) and genomic spaces, with a strong focus on gene regulatory pathways ((e.g. (Erwin & Davidson, 2002)). A clear description of ontological and epigenetic events analogous to an engineering blueprint in all these axes of information is years ahead. Moreover, it has become clear that, yet another dimension will have to be accounted for. This additional dimension is evolutionary time, and its major arena is, again, the genome. Rather than being a monotonous landscape with an occasional mutation occurring here and there, the genome is a dynamic space, frequently challenged by the insertion and/or proliferation of mobile genetic elements such as viruses and transposons (Kazazian, 2004). All these wandering elements have the potential to disrupt the genome, leading to insertion mutations, inversions, and translocations, but, more importantly, from an evolutionary standpoint, they also have the capacity to foster genome innovation, including the assembly of entirely new gene regulatory networks.

To understand the establishment and the evolution of gene regulatory networks, we have previously examined the Slow Myosin Heavy Chain III gene promoter (SMyHC III) and determined that it is able to drive preferential atrial gene expression (Bruneau et al., 2000; Nikovits et al., 1996; G F Wang et al., 1998; Gang Feng Wang et al., 1996, 2001; J Xavier-Neto et al., 1999). The sequence AGGACAAAGAGGGGA located from −801 to −787 bp upstream from the transcription start site of SMyHC III contains two Hexad sequences (Hexads A and B). Hexads A and B were previously identified as a dual putative Vitamin D Receptor Element (VDRE) and a Retinoic Acid Receptor Responsive Element (RARE) (G F Wang et al., 1998; Gang Feng Wang et al., 1996, 2001), respectively, with a well-established ventricular inhibitory function associated with the VDRE (Gang Feng Wang et al., 1996). Although correct, the current model for selective atrial expression via ventricular repression requires an extension to include atrium-specific activating elements present in the SMyHC III promoter (G F Wang et al., 1998; Gang Feng Wang et al., 1996). We sought to investigate additional mechanisms for atrial specificity in the SMyHC III promoter using *in silico* and in vivo approaches. We show that the atrial preference exhibited by the SMyHC III promoter is manifested in avian, mammals, and teleost fishes, chiefly on account of a low frequency repetitive 32 base pair genome element formed by tandem reiterations of three purine-rich hexanucleotide repeats, here designated as the complex Nuclear Receptor Element (cNRE). The cNRE is a versatile regulator of selective cardiac chamber expression, switching from SMyHC III activator to repressor functions according to atrium, or ventricular contexts, respectively. We demonstrated that the combination of three Hexads A, B, and C within the cNRE, provides an information processing platform that integrates signals from different elements and that the cNRE is necessary and sufficient to confer preferential atrial expression. Finally, using comparative genomics, we provide evidence that the cNRE was associated with the SMyHC III gene through infection of an ancestral host germline by an unknown virus resulting in recombination into the genome of a Galliform bird ancestor at the root of the Galliformes radiation in the Cretaceous, about 63 million years ago.

## Results

### Identification of the cNRE as a tripartite structure with Hexads A, B, and C

To investigate additional mechanisms of atrial specificity, we performed computerized profiling of nuclear receptor binding sites in the SMyHC III promoter. This study predicted a novel nuclear receptor binding Hexad (Hexad C), adjacent to Hexads A and B known to act as ventricular repressor sequences (aaggacaaagaggggacaaagAGGCGGaggt at -786 to -778 bp) (G F Wang et al., 1998; Gang Feng Wang et al., 1996) (Figure 1A). The combination of these three Hexads sequences (A +B +C) was designated as the complex Nuclear Receptor Element (cNRE). We postulated that this novel tripartite nuclear receptor binding contains the minimal information necessary and sufficient for specific atrial expression in vertebrate embryos.

**Figure 1.**
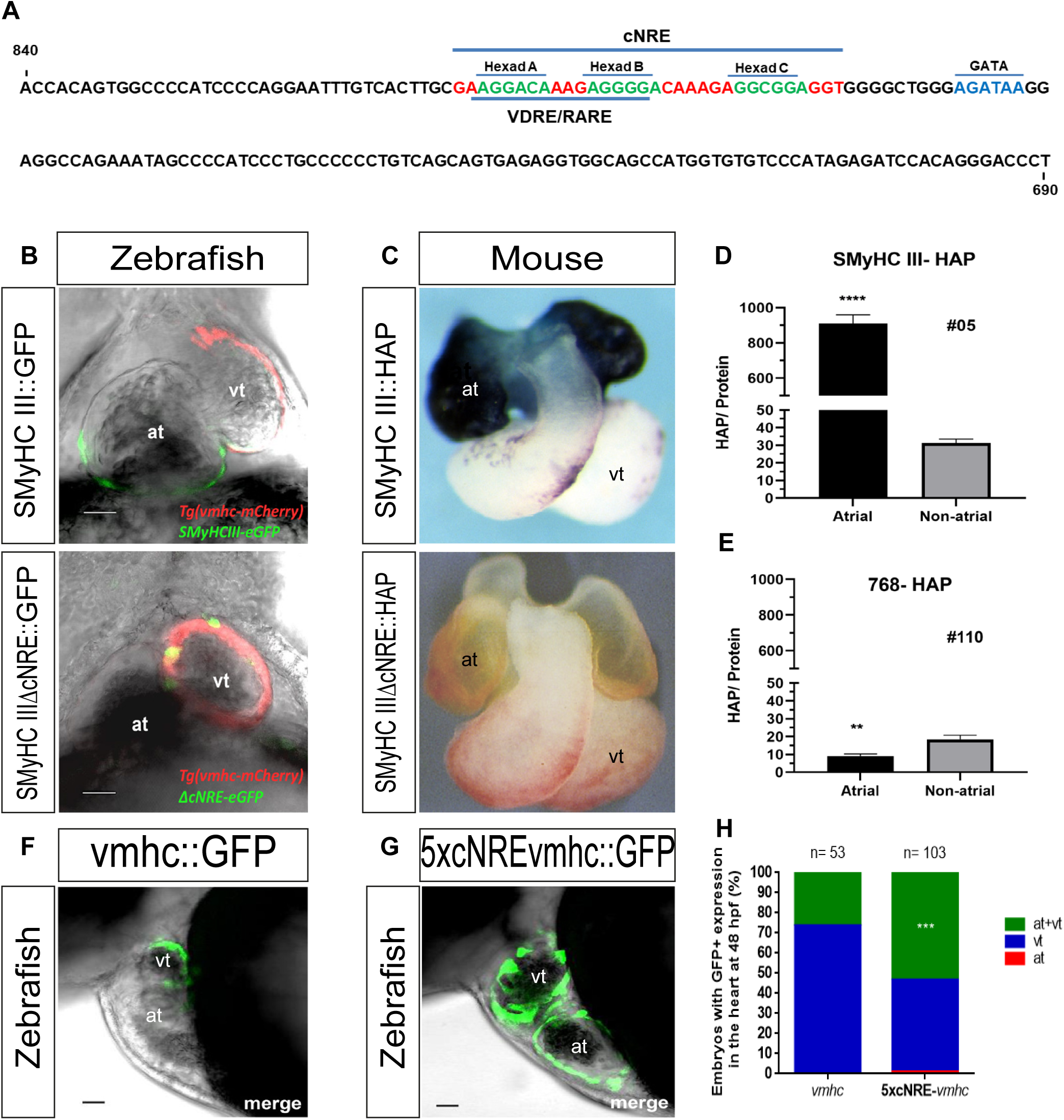
The cNRE is necessary and sufficient to drive expression in atrial cells. **A)** Schematic representation of the *SMyHC III* promoter sequence highlighting the position of the cNRE sequence. **B)** Confocal image in frontal views, anterior is to the top of a representative zebrafish embryos. Exclusive ventricular expression is demonstrated by overlapping GFP expression driven by ΔcNRE and stable mCherry fluorescence driven by the ventricular stable line. **C)** Frontal views, anterior is to the top, mouse embryos. *SMyHC III*::HAP isolated heart at 10.5 dpc showing intense, dark blue, atrial coloring indicative of high HAP expression and *SMyHC IIIΔcNRE*::HAP isolated heart at 10.5 dpc showing absence of HAP expression. **D)** HAP assays in homogenates of atrial and non-atrial cardiac tissues in *SMyHC III*::HAP (n= 18), p<0.0001. **E)** HAP assays in homogenates of atrial and non-atrial cardiac tissues in the *SMyHC IIIΔcNRE*::HAP mutant (n= 16), p=0.0013. **F)** Confocal image in lateral views, anterior is to the left of a representative zebrafish embryo. Exclusive ventricular GFP expression is observed at 48 hpf when injected with the *vmhc* promoter and **G)** a representative embryo expressing GFP in both heart chambers at 48 hpf when injected with the *5xcNRE*-*vmhc* construct. **H)** Graphical analysis of chamber expression patterns of the cohort of embryos injected with *vmhc* or *5XcNRE-vmhc* promoter constructs. (at) atrium. (vt) ventricle. chi-square test, p<0.05, comparing *vmhc::*GFP and *5xcNRE-vmhc::*GFP embryos for each condition. Scale bars are 30 µm.

### The cNRE is a transferable *cis-*regulatory agent necessary for driving atrial expression

To test the necessity of the cNRE for atrial specificity, we performed transient expression assays in zebrafish embryos (**Figure 1B**). Two reporter constructs, the quail *SMyHC III* promoter driving GFP (*SMyHC III*::GFP) and the quail *SMyHC III* promoter in which the cNRE was deleted (*SMyHC IIIΔcNRE*::GFP), were injected into the Tg(*vhmc*::mCherry) embryos (Jin et al., 2009). This line exclusively expresses mCherry in the ventricle. We assessed the proportion of embryos displaying GFP expression in only the atrium, ventricle, or both chambers (**Figure 2A, B**). With the wild-type (WT) SMyHC III promoter construct, 37% of embryos (n= 33) displayed GFP expression in the atrium, and 53% (n= 45) expressed the reporter in both ventricular and atrial chambers (**Figure 2B**). However, there was greater atrial-specific expression (37%) than ventricular-specific expression (10%, n= 6) (**Figures 1B, 2B**). In contrast, atrial-specific expression was statistically significantly reduced in the mutated quail promoter *SMyHC IIIΔcNRE*::GFP reporter assays (21%, n= 9, p=0.0026), and concomitantly, ventricular-specific expression was statistically significantly increased (38%, n= 26, p=0.0038) (**Figures 1B, 2B**). There was no statistically significant change in the proportion of embryos displaying non-chamber-specific expression for both SMyHC *III*::GFP and the *SMyHC IIIΔcNRE*::GFP constructs (53%, n= 82 and 41%, n= 55 respectively) (**Figure 2B, Figure S1**). Taking together, these results suggest that the deletion of the cNRE from the *SMyHC III* promoter reduces atrial-specificity and support the idea that the cNRE is necessary for driving atrium-specific gene expression, through the release of ventricular repression.

**Figure 2.**
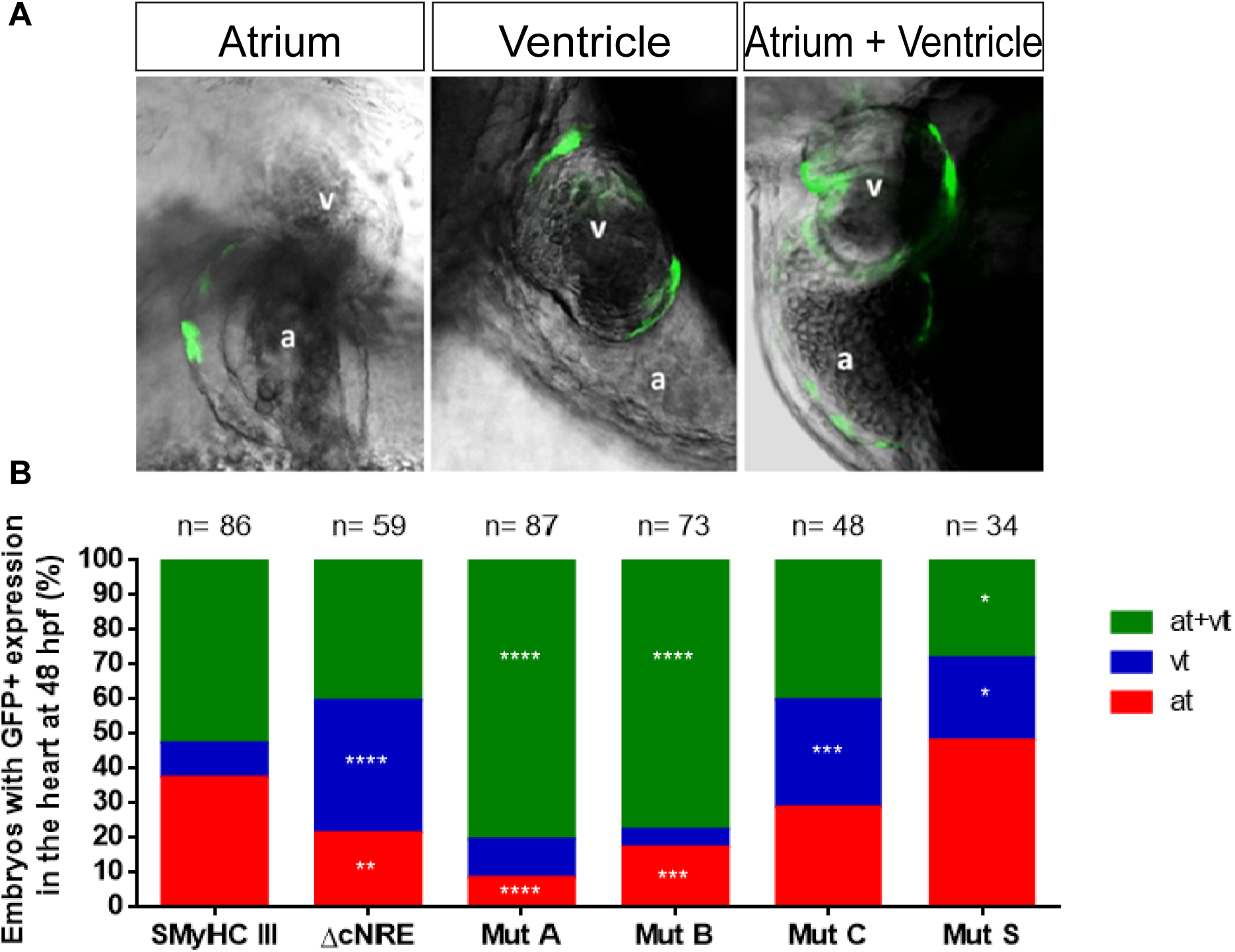
Mutational analysis of the *SMyHC III* promoter in zebrafish reveals a dual role in atrial activation and ventricular repression. **A)** Representative panel of GFP expression patterns in cardiac chambers of zebrafish embryos in lateral views, anterior to the left, injected with *SMyHC III*::GFP. **B)** Graphic representation of GFP chamber expression patterns of cohorts of embryos injected with *SMyHC III*::GFP, *SMyHC IIIΔcNRE*::GFP and constructs containing point mutations in the cNRE Hexads A, B and C (Mut A, B and C, respectively) as well as a non-Hexad control mutation (Mut S). Embryos were analyzed at 48 hpf and classified into three categories of cardiac expression patterns: exclusive atrium (at), exclusive ventricular (vt) and atrium plus ventricular (at+vt). chi-square test, p<0.05, comparing *SMyHC III* to each mutation and condition.

To support this interpretation, we performed transgenic assays in the *SMyHC III*::HAP transgenic mouse line driving atrial-specific expression of the human alkaline phosphatase reporter gene (HAP) (J Xavier-Neto et al., 1999) under the control of the quail *SMyHC III* promoter (**Figure 1C, Figure S2**). We observed that the deletion of a 72bp region encompassing the cNRE (*SMyHC III*ΔcNRE::HAP) abrogated atrial expression (**Figure 1C, Figure S3**), further supporting the notion that the cNRE is necessary for driving atrium-specific expression. A statistically insignificant level of reporter expression was observed in non-atrial regions in both the *SMyHC III*::HAP and the *SMyHC III*ΔcNRE::HAP transgenic mouse lines (**Figure 1D-E**).

### The cNRE is a transferable *cis-*regulatory agent sufficient to drive atrial expression

To test the sufficiency of the cNRE for driving atrial specific expression, we devised a conversion assay, which aimed at testing whether the cNRE is sufficient to revert a pattern from ventricular to atrial activation. To do so, we performed transient expression assays with the *vmhc* promoter driving GFP expression (*vmhc*::GFP) (**Figure 1F**) and with a 5’ fusion of five tandem repeats of the cNRE to this *vmhc* reporter construct (*5xcNRE-vmhc*::GFP) (**Figure 1G**). In 48 hpf zebrafish, we observed ventricule-specific GFP expression in most transients injected with *vmhc*::GFP (73%, n= 39) (**Figure 1H**). There were no embryos expressing GFP exclusively in the atrium. In contrast to the WT *vmhc* promoter, with the *5xcNRE-vmhc* fusion construct (**Figure 1G**), we observed a significant increase in the proportion of embryos showing both atrial and ventricular GFP expression (n= 55), a decrease in the number of embryos expressing GFP exclusively in the ventricle (46% of the embryos, n= 47), and strikingly, a single embryo with reporter expression exclusively in the atrium (0,97%, n= 1) (**Figure 1H**). These experiments demonstrated that, outside of its native context in the SMyHC III promoter, the cNRE is sufficient to convert ventricular to atrial expression.

Taking together, our results suggest that the cNRE is necessary and sufficient for directing preferential atrial gene expression. Experiments in zebrafish suggest that atrial specificity could be achieved through ventricular gene repression. Hence, we sought to investigate the precise contribution of the binding elements within the cNRE to the activation of atrial-specific gene expression or repression of ventricular-specific gene expression.

### Combinatorial recruitment of Hexads A, B and C is essential for cNRE activity *in vivo*

To understand the *cis*-regulatory composition of the cNRE we assessed the contribution of Hexads A, B and C in driving atrial specific expression *in vivo*. We used site-directed mutagenesis of individual Hexads in the *SMyHC* III promoter used in zebrafish transient assays (*SMyHC* III*::GFP*) (**Figure 2**) and in mouse mutant lines (*SMyHC* III*::HAP*) (**Figure 3, Figures S2, S3, S4, S5**). Reporter expression driven by the mutated cNRE was compared to WT cNRE (**Figure 4A**), or its complete deletion (**Figure 4B**). Mutation of Hexad A (Mut A) was obtained by substituting the Hexad A sequence 5’ AGGACA 3’ for 5’ GTCGAC 3’ (**Figure 4C**). Dinucleotide substitutions were performed on Hexad B and Hexad C to obtain Mut B and Mut C, respectively (**Figure 4D-E**). One dinucleotide substitution in the spacer region between Hexads B and C was designed as a non-Hexad control mutation (Mut S) (**Figure. 4F**). In zebrafish embryos, when comparing Mut A to the WT *SMyHC* III*::GFP* promoter (**Figures 2, 4A, A’**), we observed a statistically significant increase in the proportion of embryos with GFP positive cells in both atrium and ventricle (from 53%, n=45, in WT to 81%, n=70, in Mut A, p<0.001) and a statistically significant decrease in the proportion of embryos with GFP-positive cells exclusively in the atrium (from 38%, n=33, in WT to 8%, n=7, in Mut A, p<0.0001) (**Figure 2B**). Mut A thus leads to a significant increase of GFP-positive cells in both atrium and ventricle, mainly by stimulating GFP expression in the ventricle (**Figures 2B, 4C, C’**). We hence conclude that, in zebrafish, Hexad A is required for ventricular repression. In contrast, in mouse embryos, Mut A (**Figures 3A, 4C’’, Figure S4**) had no effect on HAP expression when compared to WT *SMyHC* III::HAP (**Figures 1C, 4A’’, Figure S2**). Similar to the effect of Mut A, for Mut B in zebrafish, we observed a statistically significant increase in the proportion of GFP-positive cells in both chambers (from 53%, n=45, in WT to 78%, n=62, in Mut B, p<0.0001) and a statistically significant decrease in the proportion of embryos with GFP-positive cells exclusively in the atrium (from 37%, n=33, in WT to 17%, n=9, in Mut B, p=0.0002) (**Figure 2B**). For Mut B, we found a similar effect in mouse embryos, with conspicuous HAP expression in the left ventricle and proximal outflow tract (**Figure 3C-H**). Altogether, the experiments in both the zebrafish and the mouse support the notion that Mut B released ventricular repression (**Figure 4D, D’, D’’**), thus suggesting that Hexad B is required for repression of gene expression in the ventricle. Mutation of Hexad C, Mut C, resulted in a statistically significant increase of ventricular-specific expression (31%, n=16, p=0.0005) when compared to the *SMyHC III* control (10%, n=6) in zebrafish embryos (**Figure 2B**). This increase is accompanied by a decrease of the proportion of embryos with GFP expression only in the atrium or in both the atrial and ventricular chambers. Consistent with this result in zebrafish, we found a marked decrease of HAP staining in mouse Mut C mutants (**Figure 3I, J, M, Figure S5B-D**), relative to WT mice (**Figure 3K, Figure S5A**). Of note, compared to the WT, Mut C mutant mice were characterized by a general reduction of HAP expression in 10.5 dpc hearts (**Figure 3L, N, Figure S5A**). Taken together, the reduction of atrial reporter expression of Mut C in both zebrafish and mouse (**Figures 4E, E’, E’’**) suggests that Hexad C acts as an atrial activator. As control of the Hexad deletions, we assessed the effect of a mutation in a spacer region outside the Hexads (Mut S). This mutation is located between Hexads B and C, adjacent to Hexad B (**Figure 4F**). In zebrafish, Mut S did not trigger a significant change in atrial-specific expression. However, we observed an increase of GFP-positive cells in the ventricle (29%, n=8) compared to the *SMyHC III* control (10%, n=6), at the expense of GFP expression in both chambers (29%, n=10, p=0.0388) compared to the *SMyHC III* control (53%, n=45) (**Figure 2B**). This suggests that Mut S might contribute to the repression of atrial expression (**Figures 4F, F’**). However, these results could not be confirmed in mice, where Mut S had no effect on HAP expression compared to the WT *SMyHC III*::HAP (**Figures 3O-P, 4F’’**). In conclusion, the phenotype displayed by Mut A and Mut B transgenic embryos is consistent with the release of a highly stereotyped ventricular expression pattern. These observations are consistent with those previously described after the deletion of the whole VDRE/RARE motif (Hexad A and B) (G F Wang et al., 1998; Gang Feng Wang et al., 1996). We further reveal that the Hexad C element in the *SMyHC III* promoter is responsible for atrial-specific activation **(Figure 4**).

**Figure 3.**
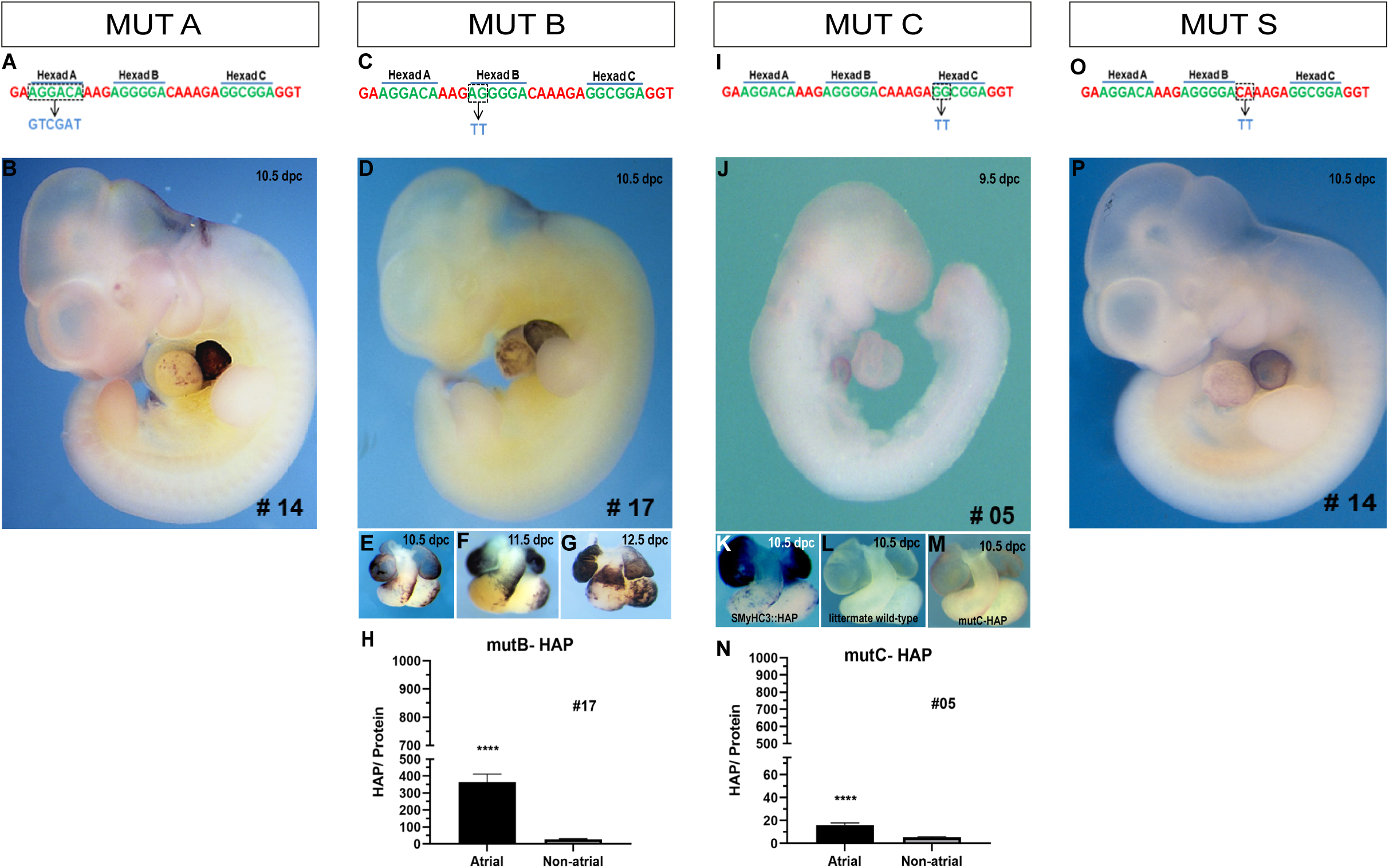
Point mutations in Hexads B and C affect HAP expression in the heart. **A)** Strategy for the mutation of Hexad A (Mut A). **B)** Mouse line 14 (Mut A) at 10.5 dpc, showing atrial-specific expression of HAP. **C)** Strategy for the mutation of Hexad B (Mut B). **D)** Mouse line 17 (Mut B) at 10.5 dpc, showing expression of HAP. **E-G)** Time course (10.5 dpc to 12.5 dpc) of cardiac expression in both chambers (atrium and ventricle) in mouse line 17 (Mut B). **H)** Comparison of HAP assays in homogenates of atrial and non-atrial cardiac tissues from line 17 (Mut B) in 10.5 dpc embryos (n= 38). **I)** Strategy for the mutation of Hexad C (Mut C). **J)** Mouse line 5 (Mut C) at 9.5 dpc. **K)** Isolated heart from the *SMyHC III*::HAP line at 10.5 dpc. **L)** Isolated heart from a wild-type littermate at 10.5 dpc. **M)** Isolated heart from mouse line 5 (Mut C) at 10.5 dpc. **N)** HAP assays in homogenates of atrial and non-atrial cardiac tissues from line 5 (Mut C) in 10.5 dpc embryos (n= 15). **O)** Strategy for the mutation of the non-Hexad control (Mut S) in the spacer sequence between Hexads B and C. **P)** Representative mouse line 14 (Mut S) at 10.5 dpc, showing atrial-specific HAP expression.

**Figure 4.**
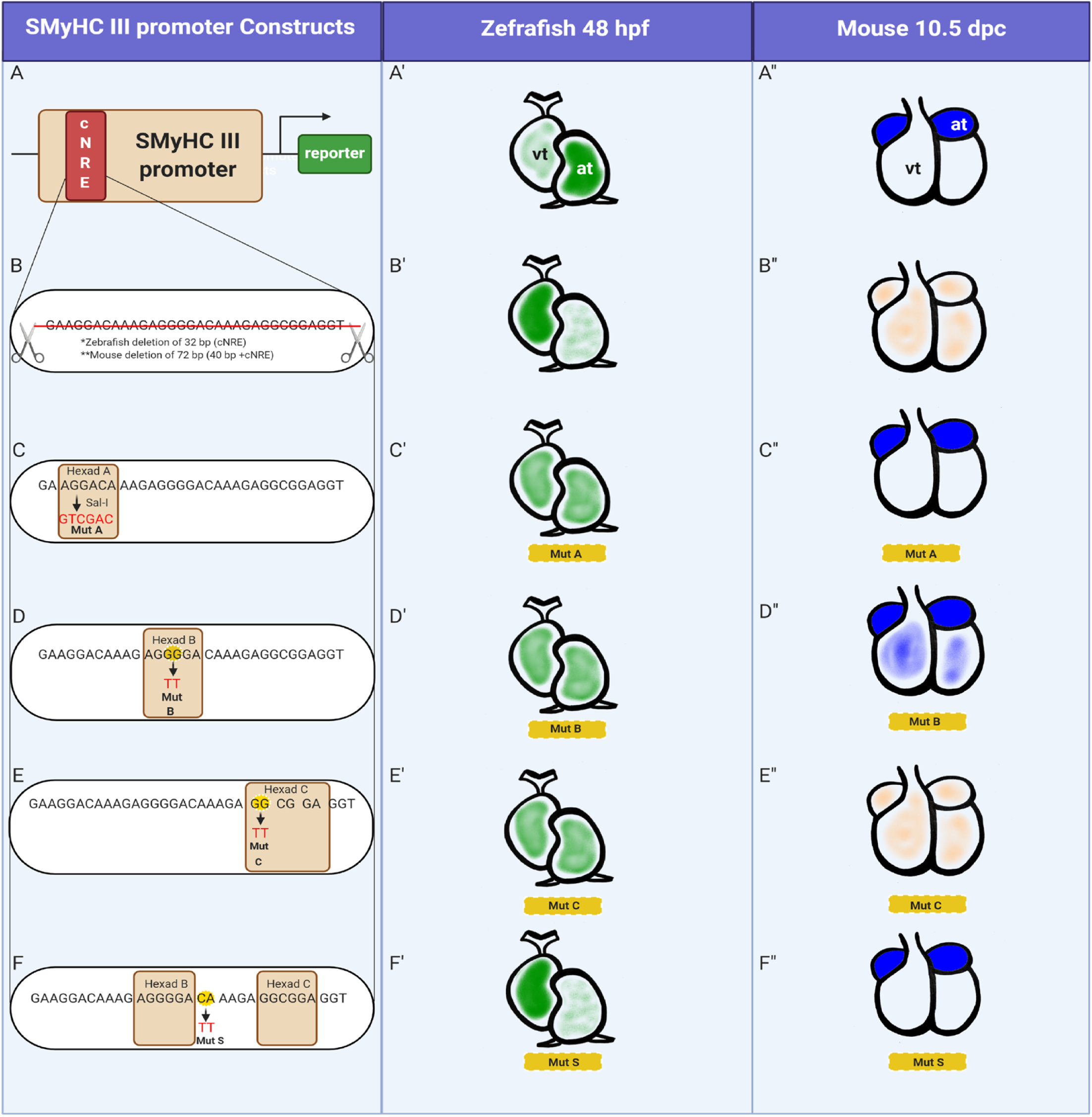
Comparison of cNRE mutations between zebrafish and mice. **A)** *SMyHC III* promoter in **A’)** zebrafish and **A’’)** mice. **B)** cNRE deletion in **B’)** zebrafish and **B’’)** mice. **C)** Mutation of Hexad A (Mut A) in **C’)** zebrafish and **C’’)** mice. **D)** Mutation of Hexad B in **D’)** zebrafish and **D’’)** mice. **E)** Mutation of Hexad C in **E’)** zebrafish and **E’’)** mice. **F)** Mutation of the spacer sequence between Hexads B and C (Mut S) in **F’)** zebrafish and **F’’)** mice. Zebrafish hearts in lateral views and mouse hearts in frontal view. (at) atrium. (vt) ventricle.

### Evolutionary origin of the cNRE

Given that the quail cNRE sequence drives preferential atrial expression in different vertebrate taxa, including zebrafish and mice, we sought to define the evolutionary origin of the cNRE. Based on the observation that the *SMyHC III* gene is a representative of a clade of slow myosin’s specific to birds (Chen et al., 1997; Nikovits et al., 1996) (**Figure 5A**). In all galliform birds, at least one hit of the 30-32 bp-long cNRE was found with a significant E-value (**Figure 5A**). The cNRE described here, composed of the three Hexads A, B and C, is thus unique to galliform birds and has likely originated in their last common ancestor, approximately 63 million years ago (Kuhl et al., 2021). Interestingly, the cNRE sequence strongly matches more than one locus in the genomes of various galliform birds, including Japanese quail (*Coturnix japonica*), chicken (*Gallus gallus*), pinnated grouse (*Tympanuchus cupido*), wild turkey (*Meleagris gallopavo*), helmeted guineafowl (*Numida meleagris*), Indian peafowl (*Pavo cristatus*), common pheasant (*Phasianus colchicus*) and Mikado pheasant (*Syrmaticus mikado*) (**Table 1**). Since galliform birds are known to be highly susceptible to viral integration (Holmes, 2011; Kaleta, 1990; McGeoch et al., 2000; Morissette & Flamand, 2010; Nair, 2005), we hypothesized that the cNRE might have a viral origin. To test this hypothesis, we systematically screened for cNRE sequences in the avian viral database (see materials and methods). We found statistically significant cNRE matches in the genomes of papillomaviruses, paramyxoviruses, retroviruses and herpesviruses (**Figure 5B**). This suggests that the cNRE might have originally been associated with the *SMyHC III* gene through viral infection of an ancestral galliform host and that integration into the genome occurred early during galliform radiation in the Cretaceous period, about 63 million years ago. Taken together, our study demonstrates that the cNRE is a tripartite *cis*-regulatory element that confers cardiac chamber specificity and arose during early diversification of galliform birds by genomic integration of a virus-derived sequence.

**Figure 5.**
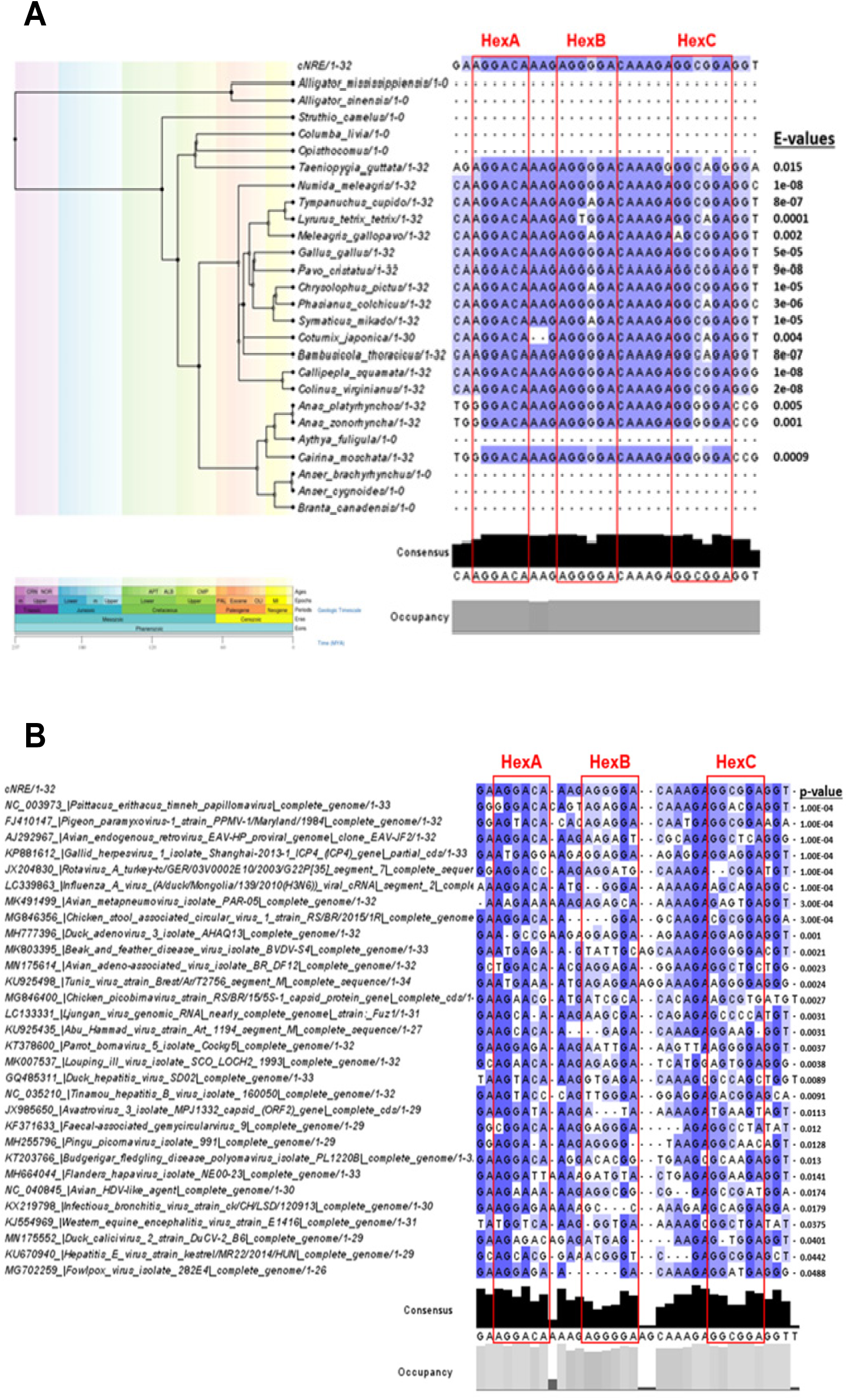
Evolutionary conservation of the cNRE. **A)** Phylogenetic tree of cNRE sequences in American alligator (*Alligator mississippiensis*), Chinese alligator (*Alligator sinensis*), ostrich (*Struthio camelus*), pigeon (*Columba livia*), hoatzin (*Opisthocomus hoazin*), zebra finch (*Taeniopygia guttata*), helmeted guineafowl (*Numida meleagris*), pinnated grouse (*Tympanuchus cupido*), black grouse (*Lyrurus tetrix*), wild turkey (*Meleagris gallopavo*), chicken (*Gallus gallus*), Indian peafowl (*Pavo cristatus*), golden pheasant (*Chrysolophus pictus*), common pheasant (*Phasianus colchicus*), Mikado pheasant (*Syrmaticus mikado*), Japanese quail (*Coturnix japonica*), Chinese bamboo partridge (*Bambusicola thoracicus*), scaled quail (*Callipepla squamata*), bobwhite (*Colinus virginianus*), mallard (*Anas platyrhynchos*), eastern spot-billed duck (*Anas zonorhyncha*), tufted duck (*Aythya fuligula*), Muscovy duck (*Cairina moschata*), pink-footed goose (*Anser brachyrhynchus*), swan goose (*Anser cygnoides*) and Canada goose (*Branta canadensis*). For each species, the sequence displaying the statistically most significant hit with the original cNRE is shown, along with the corresponding E-value. **B)** Alignment of the original cNRE with sequences identified in different viruses, along with their respective p-values. (Hex) Hexads.

**Table 1.**
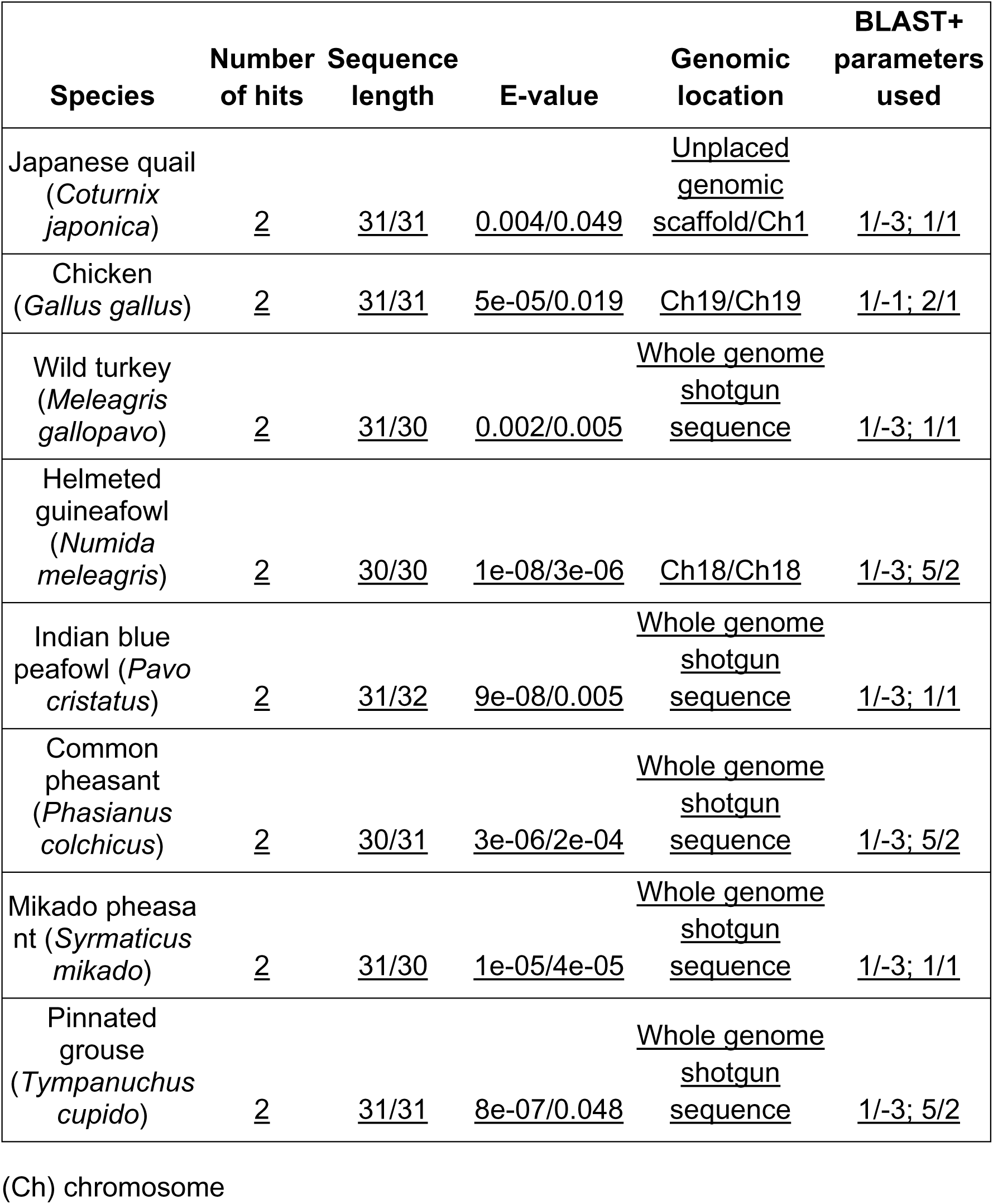
Species with multiple cNRE hits

## Discussion

Cardiac development in mammals, *i.e.*, the formation of a four-chambered heart, is characterized by a series of complex morphogenetic movements and is controlled by an intricate gene regulatory network (Waardenberg et al., 2014). In this work, we have identified the complex Nuclear Receptors Element (cNRE), a new, 32bp-long regulatory element located within the quail Slow Myosin Heavy Chain III (*SMyHC* III) promoter. We further showed that the cNRE is necessary and sufficient for driving reporter gene expression specifically in atrial cells of mice and preferentially in atrial cells of zebrafish. We also demonstrated that this specific atrial regulation mediated by the cNRE crosses species barriers (from quail to mice and zebrafish), highlighting the importance of this new element for understanding the evolution of gene regulation underlying the specification of cardiac chambers during early embryonic development. Wang *et al*. (G F Wang et al., 1998) previously demonstrated that the VDRE/RARE element, which includes Hexads A and B of the cNRE, is responsible for ventricular repression of *SMyHC* III promoter activity in avian and murine hearts. They also identified a GATA-binding element in the *SMyHC III* promoter involved in activating expression in both the atrium and the ventricle, but they failed to identify the DNA sequence driving expression of the promoter in atrial cells. By searching for potential novel nuclear receptor binding sites, we defined the cNRE as a 3’ expansion of the initial 17-bp long, adding a third Hexad. The 32-bp cNRE thus contains three Hexads, A, B and C, and is responsible for the preferential activity of the promoter in the atrium (Nikovits et al., 1996; G F Wang et al., 1998; Gang Feng Wang et al., 1996, 2001). To functionally characterize the activity of the cNRE in the heart, we took advantage of the power of transient expression assays in zebrafish embryos as a complementary strategy to the generation of stable transgenic lines in mice (Meyers, 2018). The evaluation of the effects of point mutations in the cNRE in both zebrafish and mice ultimately allowed us to assess the role of each of the three Hexads constituting the cNRE. We found that cNRE can drive specific reporter gene expression in the atrium of both zebrafish and mice, as deletion of the cNRE within the *SMyHC III* promoter abrogated the atrial-specific expression driven by this promoter.

Of note, while the activity of the *SMyHC III* promoter is limited to the atrium in mice, the promoter drives expression in both chambers of the zebrafish heart. It might be that the teleost fish-specific whole genome duplication (REF) has secondarily altered the regulatory landscape controlling heart development (Jin et al., 2009). However, preferential atrial expression was nonetheless significantly decreased in the zebrafish ΔcNRE mutant, and the multimerized cNRE construct still clearly shifted the activity of the zebrafish *vmhc* promoter in an atrial direction. We sought to identify which regions in the cNRE sequence are critical to its activity by creating constructs containing mutations in the cNRE sequence for transient and stable transgenic analyses in, respectively, zebrafish and mice. When mutating the sequences of Hexad A and B in zebrafish, we observed a decrease in atrial expression concomitant with an increase in the number of individuals with expression in both chambers. In mice, in contrast, mutation of Hexad A did not affect expression, while the mutation of Hexad B increased expression in ventricular chambers. These data suggest that Hexad B, and potentially Hexad A, act as ventricular repressors. This is consistent with previous *in vitro* studies performed in quail atrial cells, where removal of Hexad A and B led to increased reporter gene expression in ventricular cells (Gang Feng Wang et al., 2001). Mutation of Hexad C resulted in a statistically significant increase of ventricle-specific expression in zebrafish. We hypothesize that this effect is due to the specific loss of atrial expression in embryos that would normally be characterized by transgenic activity of the construct in both chambers. This notion is consistent with our finding that mutation of Hexad C in mice led to an abrogation of atrial expression. We thus conclude that Hexad C plays an important role in atrial activation. In summary, we demonstrate that the cNRE contains information needed for both atrial activation and ventricular repression of the *SMyHC III* promoter. Further work will be required to define the transcription factors binding to the cNRE sequence to regulate its activity. Although the *SMyHC III* promoter is not conserved between species, it is capable of driving atrium-specific expression in different animal models, including chicken, mouse and zebrafish (Nikovits et al., 1996; G F Wang et al., 1998; Gang Feng Wang et al., 1996, 2001; J Xavier-Neto et al., 1999). We postulate that the cNRE acts as a dual *cis*-regulatory module integrating both activating and repressing signals, some of which likely mediated by members of the nuclear receptor superfamily via the VDRE/RARE binding sites. We propose that the cNRE *cis*-regulatory module emerged from an endogenous viral element (Holmes, 2011), a genomic remnant of a rare recombination event caused by a viral infection of an ancestral host. Using comparative genomics, we provide evidence that this virus-derived cNRE was associated with the *SMyHC III* gene in the last common ancestor germline of Galliform birds in the Cretaceous period, about 63 million years ago.

## Conclusion

Supported by *in vivo* experiments in zebrafish and mice, we demonstrate that the cNRE, a sequence of merely 32 bp, carries information to control both atrial activation and ventricular repression. We further provide evidence this *cis*-regulatory element is of viral origin. Our work thus highlights the evolution of specific regulatory motifs in a sequence that is present exclusively in avian genomes but can drive preferential atrial expression in different vertebrate taxa. Taken together, this study sheds light on the origin of enhancers and defines the minimum amount of information required for regulating gene expression in an atrial-specific fashion.

## Materials and Methods

### Bioinformatics profiling of nuclear receptor binding sites at the SMyHC III promoter

We devised a simulation approach to identify nuclear receptor binding sites (Hexads) in the SMyHC III promoter. The principle of the approach is based on the Poisson-Boltzmann theory and aims at calculating interaction energies between nuclear receptors and DNA as an approximation of their respective binding affinities. Protein/DNA complexes were assembled for molecular dynamics profiling by positioning three-dimensional (3D) structures of nuclear receptor DNA-binding domains on the 3D structure of the cNRE DNA. RXR, RAR, and VDR crystal structures are available at the Protein Data Bank (http://www.rcsb.org) with the codes 1DSZ, 1KB4 and 1BY4, respectively (Rastinejad et al., 2000; Shaffer & Gewirth, 2002; Zhao et al., 2000). The free binding energy was calculated for protein/DNA complexes using the trajectories obtained from the molecular dynamics profiling with the software MM-PBSA in the AMBER package (Sekijima et al., 2003). Each complex (AB) was split into two parts: the nuclear receptor structure (A) and the cNRE structure (B), and the energy was calculated for the whole complex AB as well as for each part, A and B. Binding energy differences were obtained according to: ΔΔG = ΔGAB - (ΔGA + ΔGB). Binding free energies for all cNRE Hexads were plotted with reference to its first nucleotide. Data were pooled and analyzed as a box plot to identify values below the 10th percentile. These values were used to identify all potential Hexads within the cNRE. To quantify the potential for nuclear receptor binding within the cNRE we scored the number of times a given cNRE nucleotide was part of a Hexad.

### Generation of mouse reporter lines

We generated mice containing mutations in critical nucleotides of Hexads A, B and C (Mut A, B and C, respectively) as well as a non-Hexad control mutation (Mut S). The constructs were synthesized with Agilent QuikChange II XL Site-Directed Mutagenesis Kit® (Cat #200521) following the manufacturer’s instructions and using the primers described in Table 2. Constructs were subsequently sequenced to confirm the substitutions. The Hexad A mutation (Mut A) was generated by mutagenesis of this Hexad’s nucleotides to a Sal I restriction enzyme site (5’-AGGACA-3’ to 5’-GTCGAC-3’). The mutation in Hexad B (Mut B) were point mutations of the two first nucleotides of this Hexad (5’-AGGGGA-3’ to 5’-TTGGGA-3’). Similarly, the mutation of Hexad C (Mut C) consisted of point mutations of the two first nucleotides of this Hexad (5’-GGCGGA-3’ to 5’-TTCGGA-3’). The non-Hexad control mutation (Mut S) was obtained by mutating two nucleotides in the putative nucleotide spacer between Hexads B and C (5’-AGGGGAcaaagaGGCGGA-3’ to 5’-AGGGGAttaagaGGCGGA-3’). *SMyHC III*::HAP transgenic lines 1 and 5 have previously been described (J Xavier-Neto et al., 1999), and four new *SMyHC III*::HAP transgenic lines were generated (2, 6, 27 and 29). Three mutB::HAP (5, 17 and 19) and seven mutC::HAP (3, 5, 7, 14, 15, 23 and 24) transgenic lines were established.

**Table 2.**
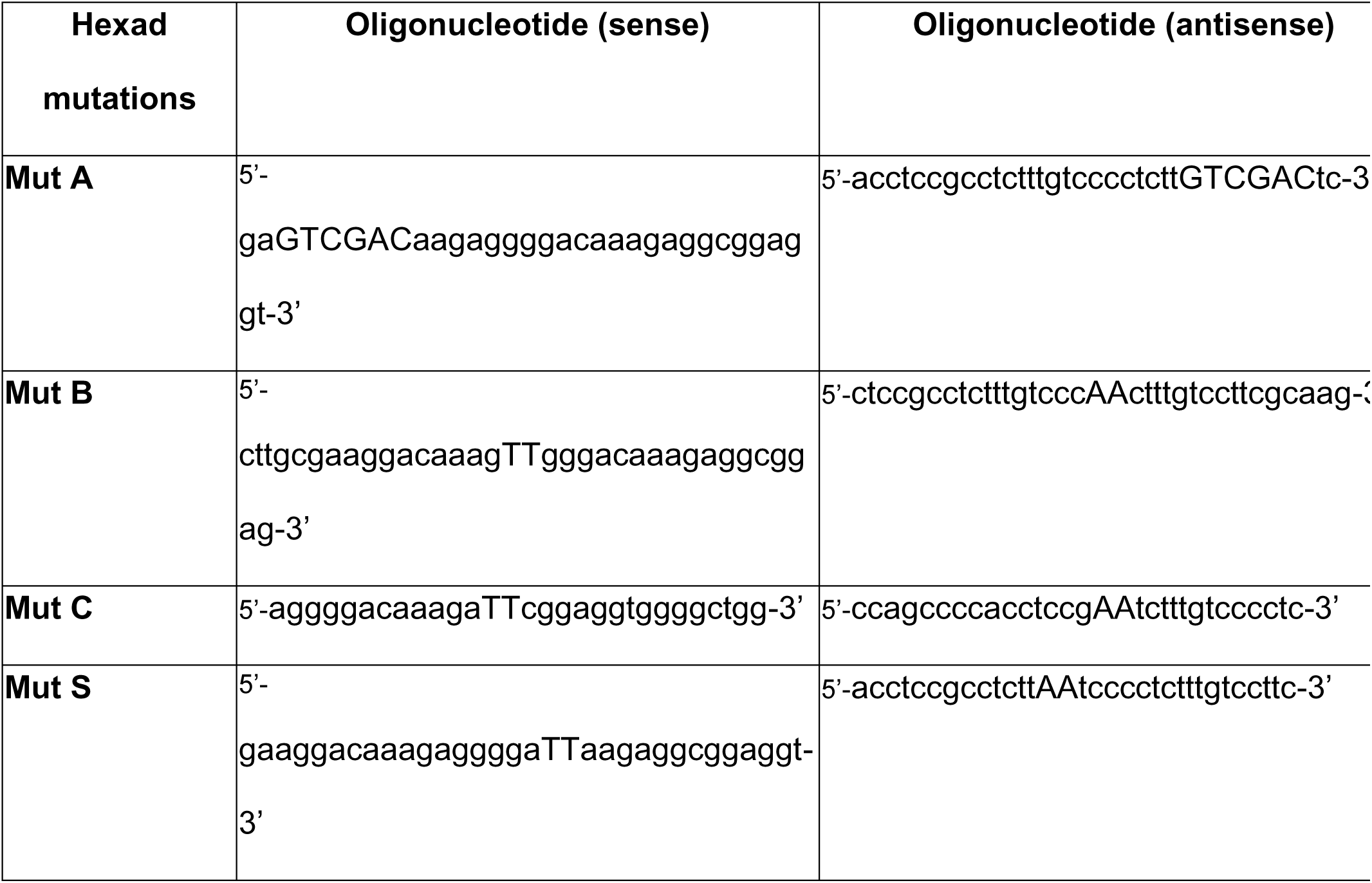
Oligonucleotides (in sense and antisense) used for Hexad mutations.

### HAP staining and histology

For HAP staining and paraffin sections, embryos and hearts were handled as described in (J Xavier-Neto et al., 1999). For HAP assays tissues were homogenized in a lysis buffer containing 0.2% Triton X-100 with a mini bouncer. HAP activity in cardiac tissues was measured with Phospha-Light^TM^, a chemiluminescent assay from PerkinElmer. HAP activity was normalized relative to protein concentration.

### Zebrafish wild-type and stable transgenic Tg(*vmhc*::mCherry) lines used in the transient experiments

To generate the Tg(*vmhc*::mCherry) line, we used a promoter fragment of 1.9 kb upstream of the *vmhc* gene, as previously described (Jin et al., 2009; Zhang & Xu, 2009). The PCR-amplified fragment was cloned into pT2AL200R150G (courtesy of Dr. Koichi Kawakami) using the Xho I and Hind III restriction sites. The eGFP, between the Cla I and BamH I restriction sites, was substituted by mCherry. Injected embryos were raised to adulthood and an F2 generation was established. To generate the *Tol2-SMyHC III*::GFP construct, the 840-bp upstream regulatory sequence of quail *SMyHC III* was excised at the Sma I and Hind III restriction sites from the *SMyHC III* pGl3 plasmid and cloned into the pT2AL200R150G using the Xho I and Hind III restriction sites. Deletion of the 32- bp cNRE from the *SMyHC III* promoter (*SMyHC III*::ΔcNRE) was obtained by digesting the pGl3 plasmid with Xho I and Hind III followed by subsequent cloning into pT2AL200R150G at the Sma I and Hind III restriction sites. For the mutational analyses of the *SMyHC III* promoter, specific mutations of the *Tol2-SMyHC III*::GFP plasmid were generated using the Agilent QuikChange II XL Site-Directed Mutagenesis Kit^®^ (Cat #200521) following the manufacturer’s instructions. Primers are listed in Table 2.

To construct the chimeric promoter *5xcNRE-vmhc*, we cloned five tandem repeats of the cNRE into the Xho I site of the *Tol2-vmhc*::GFP vector. The 5xcNRE sequence was obtained by annealing the following two oligonucleotides: 5’-CTAGGAAGGACAAAGAGGGGACAAAGAGGCGGAGGTGAAGGACAAAGAGGGGACAAAGAGGCGGAGGTGAAGGACAAAGAGGGGACAAAGAGGCGGAGCTGAAGGACAAAGAGGGGACAAAGAGGCGGAGGTGAAGGACAAAGAGGGGACAAAGAGGCGGAGGTCTCGAGA-3’ (sense) and 5’-GATCTCTCGAGACCTCCGCCTCTTTGTCCCCTCTTTGTCCTTCACCTCCGCCTCTTTGTCCCCTCTTTGTCCTTCACCTCCGCCTCTTTGTCCCCTCTTTGTCCTTCACCTCCGCCTCTTTGTCCCCTCTTTGTCCTTCACCTCCGCCTCTTTGTCCCCTCTTTGTCCTTC-3’ (antisense). For the transient transgenic assays, all *Tol2*-based constructs were co-injected (∼1nL) into the cytoplasm of one-cell stage embryos with transposase mRNA, which was transcribed from the pCS-TP vector using the mMESSAGE mMACHINE SP6 Kit (Ambion). The master mix for injections was freshly prepared with 125 ng of the plasmid of interest, 175 ng of transposase mRNA, 1 µL of 0,5% phenol red and water to complete the final volume to 5 µL (Suster et al., 2009). All constructs were microinjected in at least two independent experiments. Zebrafish embryos were staged and maintained at 28.5°C, as previously described (Westerfield, 2000), and analyzed at 48 hpf.

### Image analyses and processing

All imaging analyses were performed under a NIKON SMZ 25 fluorescent stereomicroscope, and confocal imaging was carried out using a Leica SP8 microscope.

### Statistical tests

We used either a chi-square test or an unpaired t-test (non-parametric when appropriate), and, for both analyses, a 95% confidence value was used to assess significance (p < 0.05). The data were presented in column graphs with standard deviation using GraphPad Prism 6. Each experimental group was compared to its respective control group.

### Searching animal genomes for cNRE-like sequences

To search different animal genomes for cNRE-like sequences, the command-line version of BLAST (BLAST+, version 2.5.0) was used. Animal taxa included in the analyses were those with a genome assembly available on NCBI and with phylogenetic information available on TimeTree.org. The search was conducted using blastn-short, as this allowed for a search tailored to a query sequence as short as the cNRE. All parameters were set to default, except for word size (which was always set to 4 to maximize search accuracy) and gapopen, gapextend, reward and penalty, all of which were changed according to the search parameter stringency (Table 3).

**Table 3.** Search parameters for cNRE sequence queries.

| Parameter stringency | Gapopen | Gapextend | Reward | Penalty |
| --- | --- | --- | --- | --- |
| Lenient | 2 | 1 | 1 | -1 |
| Moderate | 1 | 1 | 1 | -3 |
| Stringent | 5 | 2 | 1 | -3 |

While a gapopen/gapextend ratio of 1/-3 (in the moderate and stringent parameter sets) yielded a sequence conservation of 99%, a ratio of 1/-1 (in the lenient parameter set) resulted in a sequence conservation of 75%. Hits matching at least 30 bp of the 32-bp cNRE were retained.

Hit sequences against animal genomes were individually extracted from the databases, since BLAST+ is a local alignment algorithm that only yields parts of hit regions within a given genome. Each animal genome was thus loaded into R to obtain the exact position of the cNRE hit from the BLAST+ alignment. If the best hit for the cNRE was on the minus strand, the genome sequence was reverse complemented. Each sequence was subsequently extracted from the genome based on the genomic location of the initial BLAST+ hit and manually refined to allow a full alignment of the cNRE with a given genome sequence. Of note, the *Coturnix japonica* cNRE sequence was the only one, where gaps needed to be introduced to properly align the cNRE. The gaps were manually introduced into the extracted sequence using R to match the sequence in the BLAST+ results. All extracted sequences were compiled into a single FASTA file and aligned using CLUSTAL OMEGA. The output alignment file was processed in Jalview to create the final alignment figure.

### Searching through the viral genomes

All available avian viruses in the NCBI virus database (as of February 19, 2020) were scanned for the cNRE sequence. For this, the R function pairwise alignment (in the Biostrings package, version 2.52.0) was used to create, for each virus, a local-global alignment score of the single best hit for the cNRE sequence and the cNRE reverse complement sequence. The “pattern” was set to the viral genome, the “subject” to the cNRE (or the cNRE reverse complement), the “type” to “local-global” and all other settings were set to default. To calculate the p-value corresponding to a viral hit, 10,000 32 bp-long random DNA sequences matching the cNRE length were generated and the pairwise alignment of each sequence was determined against each viral genome. The same set of random sequences was used to determine the p-value of every virus analyzed. If a cNRE (or a cNRE reverse complement) hit in a viral genome had a local-global pairwise alignment score of more than 95% of the alignment score of the random sequences against this viral genome, this hit was considered as statistically significant (**Figure S6**). We retained top 2,000 viral genomes, based on the pairwise alignment scores of the cNRE (or cNRE reverse complement). Of these, only the best scoring virus of each viral family was chosen for further analysis. This measure ensured that no viral species or genus was overrepresented in the list of hits. Of the 2,000 viral genomes, we thus obtained 27 unique virus families and 4 unclassified viral hits. The p-value of these 31 viruses was subsequently determined, with viruses returning a p-value of 0.05 or less having been included in **Figure 5**. These hit sequences of viral genomes were saved in a single FASTA file and the FASTA file was processed using CLUSTAL OMEGA to produce a multiple sequence alignment. If a hit in a virus genome was to the cNRE reverse complement, the viral hit sequence was reverse complemented before it was included in the FASTA file. The output file was then opened in Jalview and processed to highlight different levels of sequence conservation.

## Acknowledgments

We thank Dr. Koichi Kawakami for providing the plasmid used to produce transgenic zebrafish. We thank the members of the Ramialison group and Akriti Varshney, Gulrez Chahal, and Julian Stolper for their feedback and support and Jeannette Hallab, Jeanette Rientjes, Ekaterina Salimova for their assistance with experimental troubleshooting and design. We thank the members of the Monash Bioinformatics Platform for invaluable advice, especially Stuart Archer, Adele Barugahare, Paul Harrison, David Powell, Michael See and Nick Wong. We thank the ARMI FishCore staff and Melbourne University Fish Facility. We also thank Lucas Buscaratti for his drawing created with BioRender.com that helped us to illustrate the different mutations on the *SMyHC III* promoter comparing mice and zebrafish.

## Declarations

### · Ethics approval and consent to participate

We confirm that all relevant ethical guidelines have been followed, and any necessary IRB and/or ethics committee approvals have been obtained. The present study was approved by Ethics Committee on Animal Use, Institute of Biomedical Sciences, University of São Paulo (CEUA-ICB/USP), by Ethics Committee on Animal Use, Brazilian Center for Research in Energy and Materials (CEUA-CNPEM) protocol number 24, by the University of Melbourne guidelines and local ethics committee and by the Monash University Animal Ethics Committee.

### · Availability of data and materials

The datasets used and/or analyzed during the current study are available from the corresponding authors on reasonable request, and available in the GitHub repository: https://github.com/mart-nik/cNRE_project.git.

### · Authors’ contributions

LNS generated transgenic zebrafish, contributed to discussions, and wrote this manuscript. MN and MR performed bioinformatics analyses, contributed to discussions and co-wrote the manuscript. AMSC, PJ and SD contributed to transgenic analyses. HAC, MV, TGFM and ACS carried out experiments on transgenic mice. MBG and PO performed bioinformatics analyses. FES, NR, GFW, WK and MS contributed to the conception of this study. JXN was responsible for the project design, mouse transgenics, scientific discussions and manuscript writing. All authors have read and approved the final manuscript.

---

**Supplementary Figure 1.**
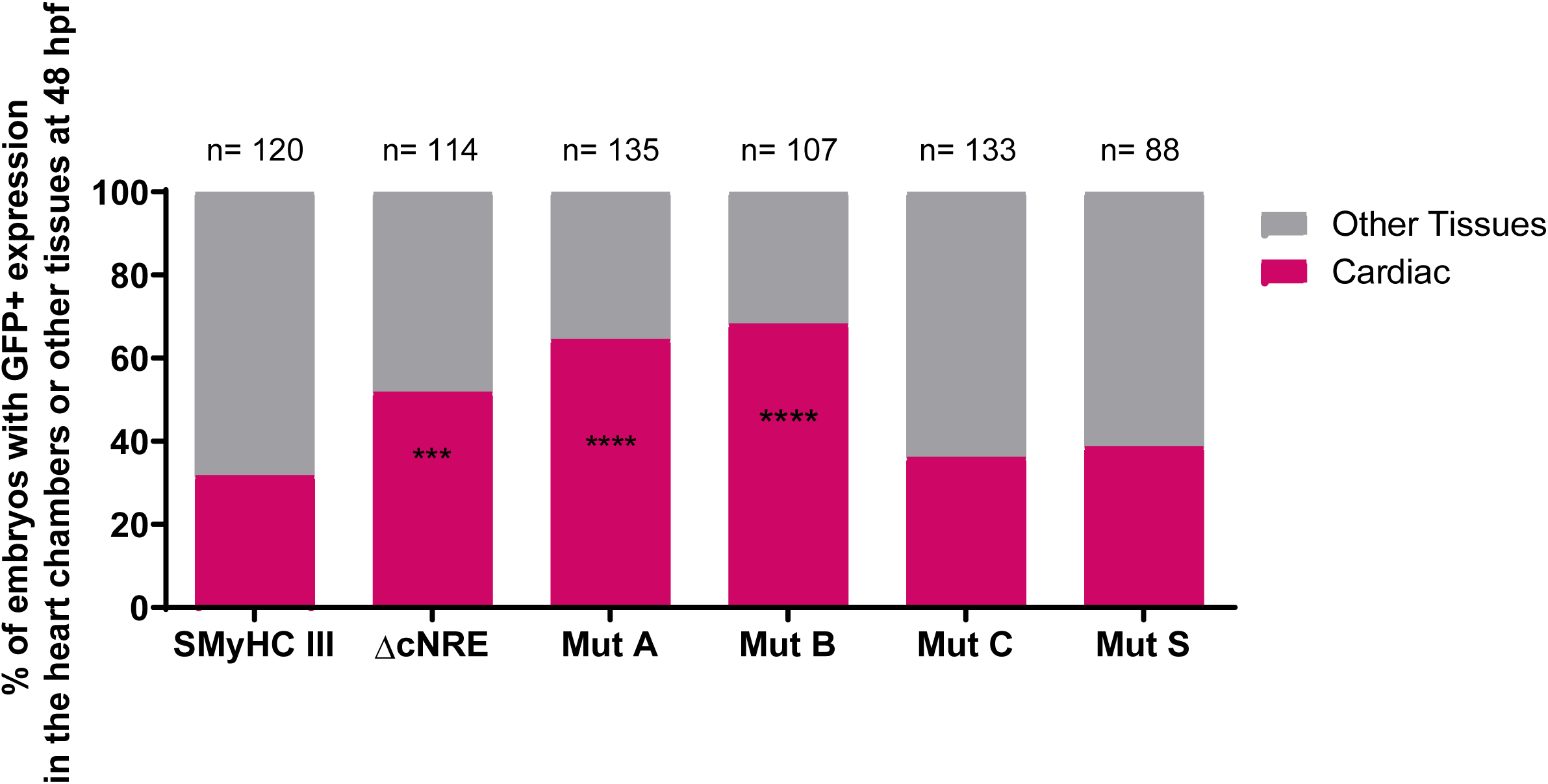
GFP expression in cardiac and non-cardiac tissues of zebrafish embryos at 48 hpf following injection with different constructs of the *SMyHC III* promoter. Graphic representation of GFP expression in cardiac and other tissues of cohorts of embryos injected with different *SMyHC III* promoter constructs and analyzed at 48 hpf. chi-square test, p<0.05, comparing *SMyHC III* to each mutation and condition.

**Supplementary Figure 2.**
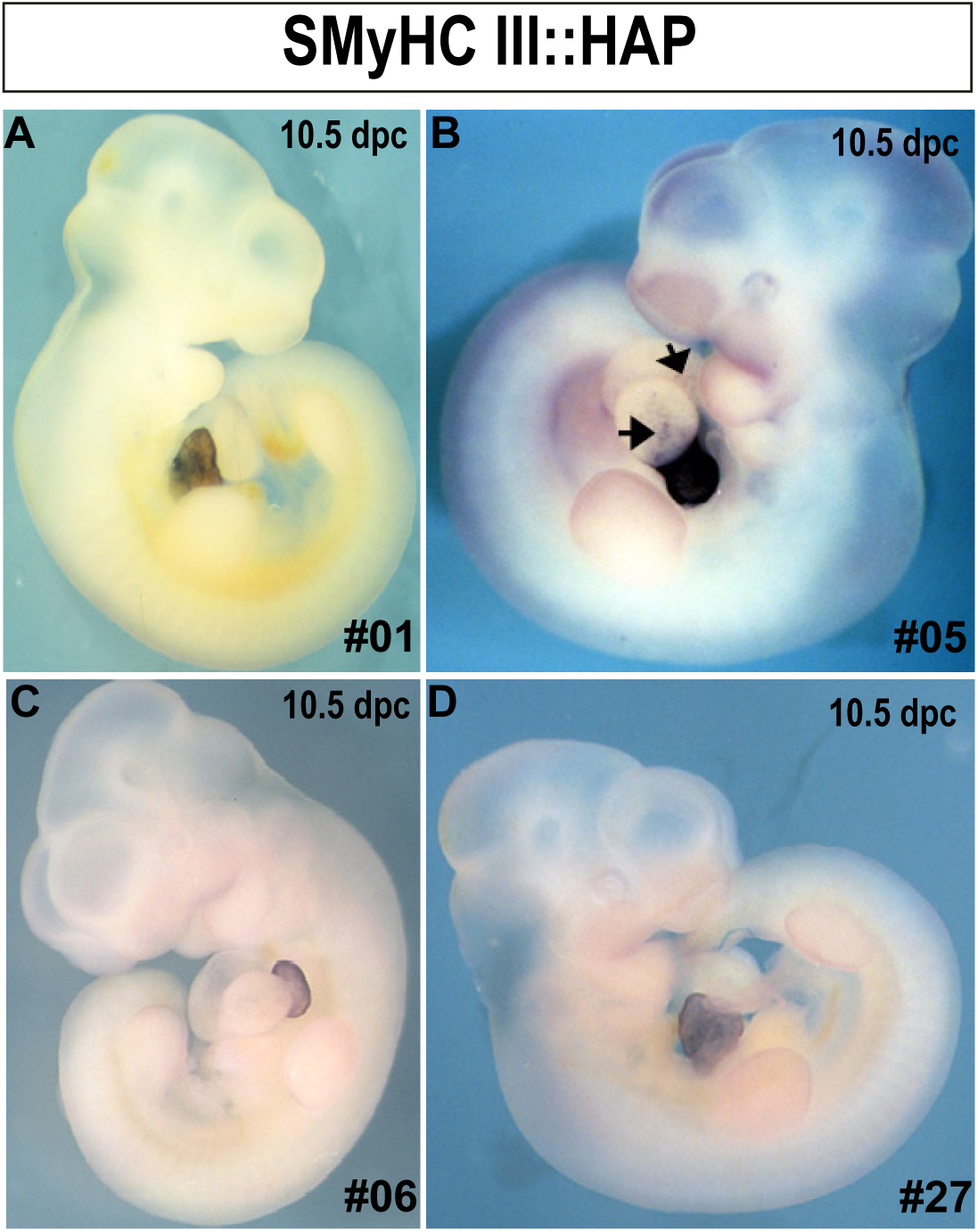
*SMyHC III*::HAP embryos expressing HAP specifically in the atrium. A-D) HAP expression at 10.5 dpc in mouse lines 1, 5, 6 and 27.

**Supplementary Figure 3.**
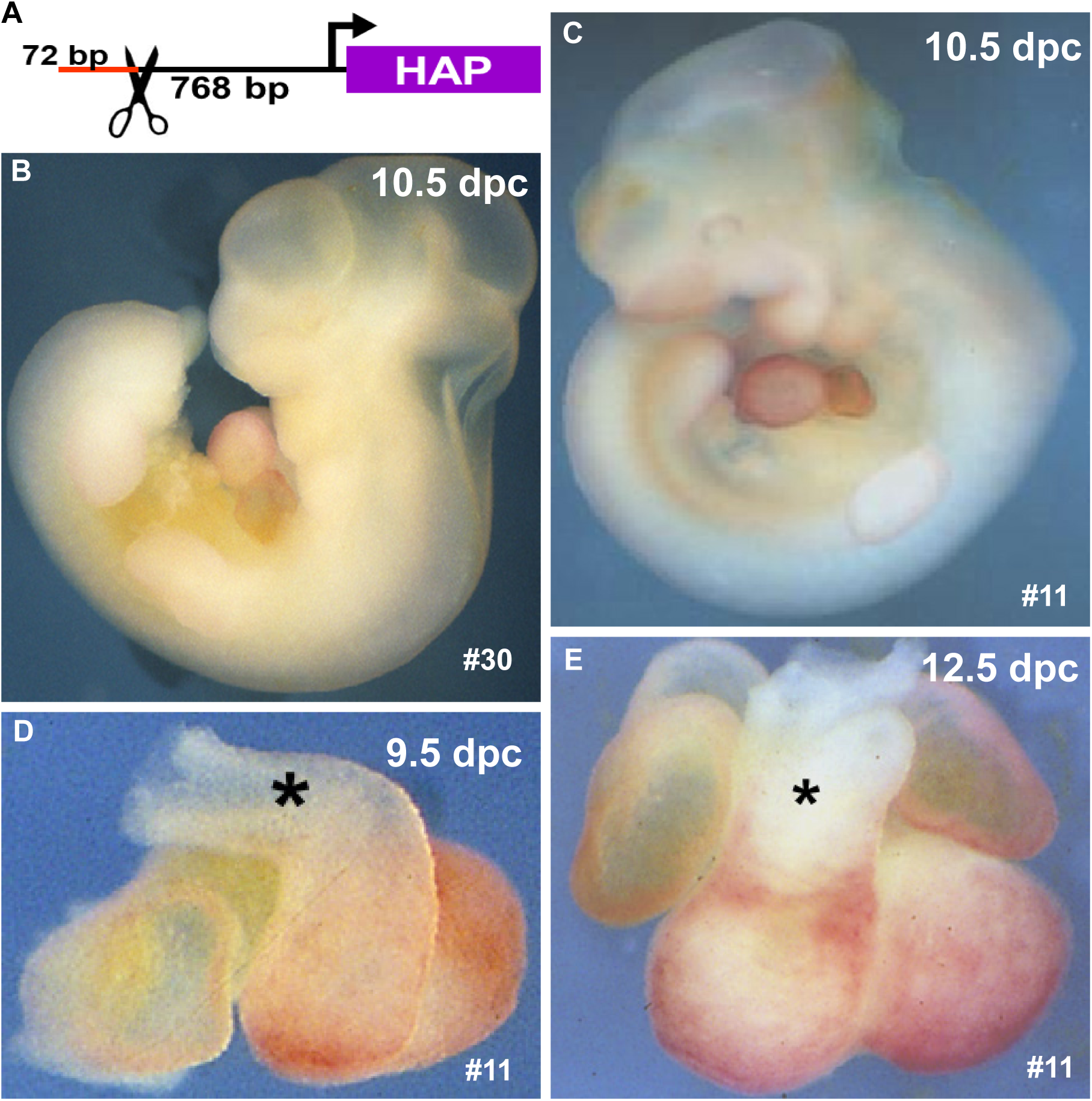
Mutation of the distal 72 bp fragment containing the cNRE sequence alters atrial HAP expression driven by the *SMyHC III* promoter. **A)** Strategy for the deletion of the distal 72 bp of the *SMyHC III* promoter. **B)** *SMyHC IIIΔcNRE*::HAP at 10.5 dpc in mouse line 30. **C)** *SMyHC IIIΔcNRE*::HAP at 10.5 dpc in mouse line 11. **D)** Isolated embryonic heart of *SMyHC IIIΔcNRE*::HAP mouse line 11 at 9.5 dpc. **E)** Isolated embryonic heart of *SMyHC IIIΔcNRE*::HAP mouse line 11 at 12.5 dpc. (*) distal outflow.

**Supplementary Figure 4.**
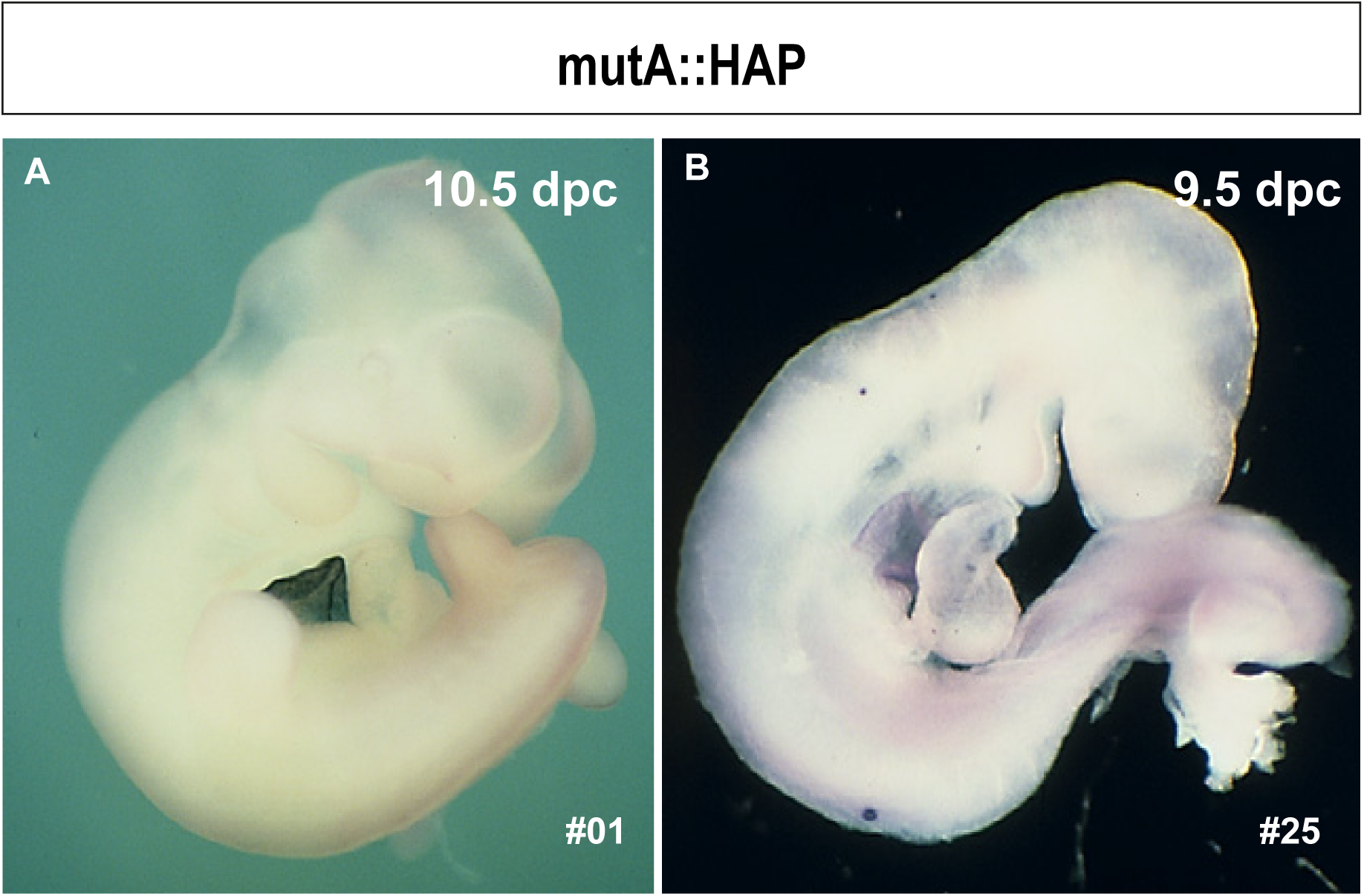
Mutation of Hexad A does not affect atrial specificity of the *SMyHC III* promoter. **A)** Mouse line 1 with mutation of Hexad A (*mutA*::HAP) at 10.5 dpc. **B)** Mouse line 25 with mutation of Hexad A (*mutA:*:HAP) at 9.5 dpc.

**Supplementary Figure 5.**
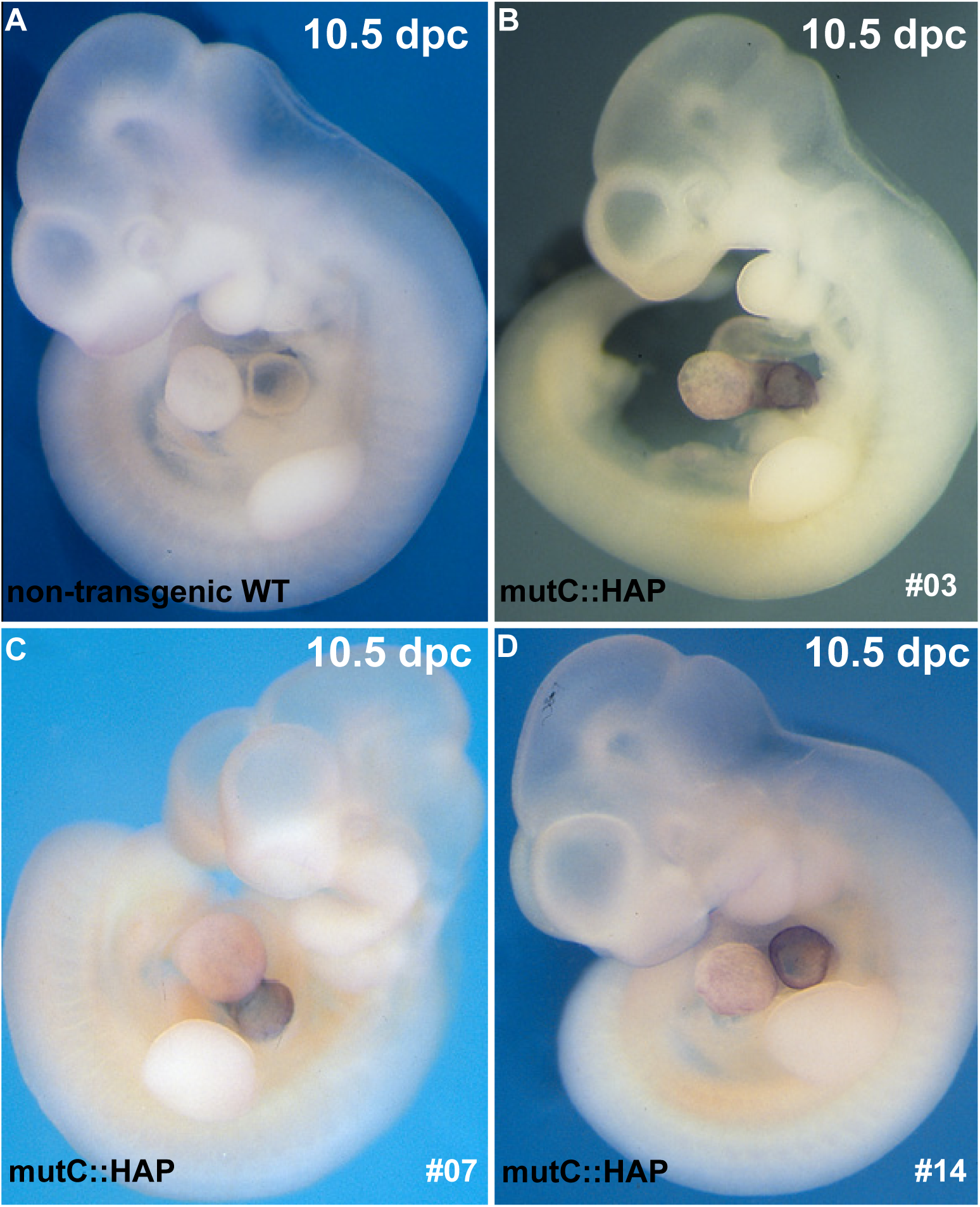
Mutation of Hexad C affects both atrial and ventricular expression of the *SMyHC III* promoter. **A)** A wild-type non-transgenic embryo displaying background HAP staining at 10.5 dpc. **B)** Mouse line 3 with mutation of Hexad C (*mutC*::HAP) at 10.5 dpc. **C)** Mouse line 7 with mutation of Hexad C (*mutC*::HAP) at 10.5 dpc. **D)** Mouse line 14 with mutation of Hexad C (*mutC*::HAP) at 10.5 dpc.

**Supplementary Figure 6.**
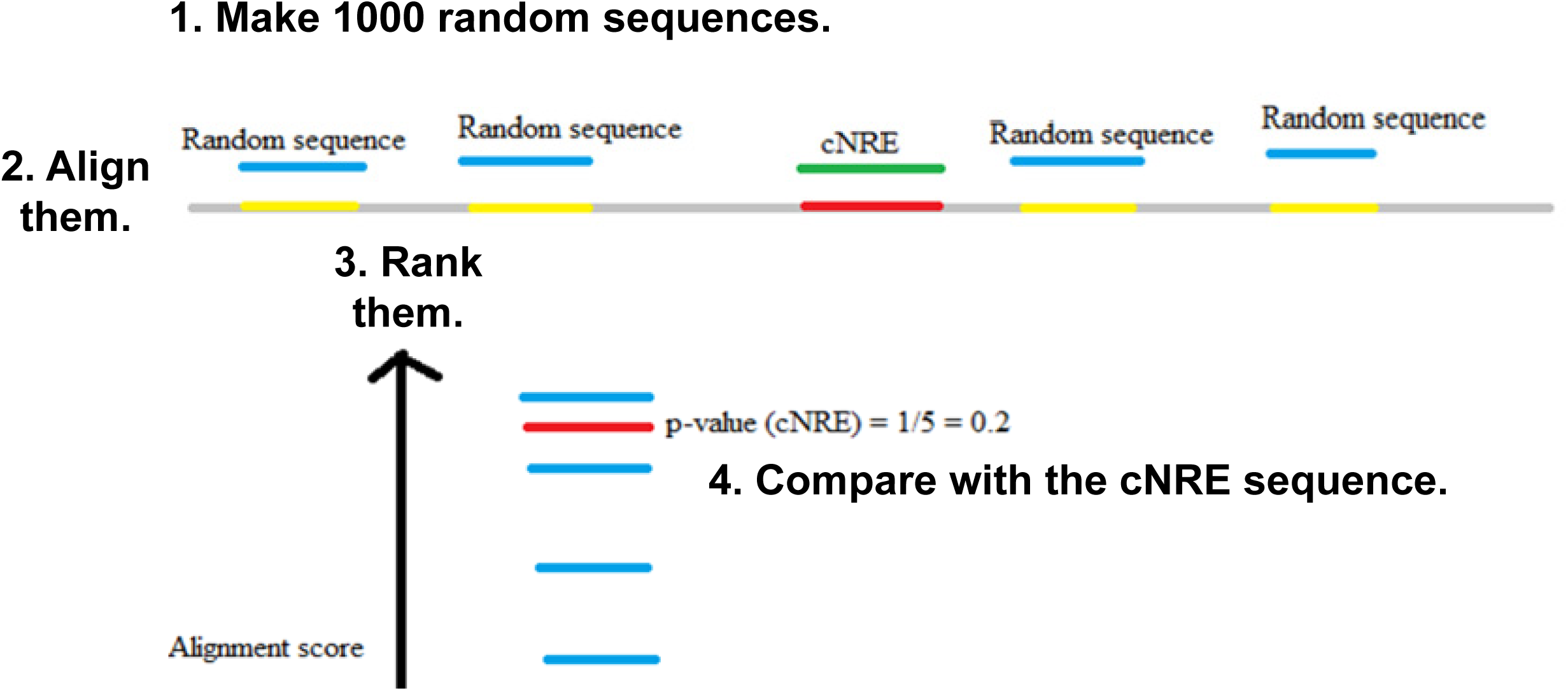
The R-based pairwise alignment method for calculating p-values for viral hits. 1000 random sequences were produced and their pairwise alignment scores were compared to those obtained with the cNRE. A viral hit was considered as statistically significant, when the pairwise alignment score of the cNRE sequence was more than 95% higher than the score obtained with the random sequences.

